# KymoMerge: a new tool for analysis of multichannel kymographs

**DOI:** 10.1101/2021.11.29.470387

**Authors:** Lloyd P. McMahon, Laura Digilio, Alois Duston, Chan Choo Yap, Bettina Winckler

**Affiliations:** Department of Cell Biology, University of Virginia, 1340 Jefferson Park Avenue, Pinn Hall 3226, Charlottesville, VA 22908, USA

## Abstract

Kymograph analysis is commonly used by researchers to study dynamic processes in cells. Current tools widely available only allow for analysis of one channel kymographs. Here we provide a python-coded, open source program to use as a plug-in for ImageJ which creates one kymograph from the kymographs of two separate channels of one time-lapse movie. This add-in, which we call KymoMerge, therefore allows for analysis of only the co-located tracks in multi-channel movies.

## Introduction

Kymographs are a commonly used method of tracking dynamic processes in cells. The method translates time-dependent microscope data (time-lapse movies) into a two-dimensional representation showing position versus time-generally with position on the x-axis and time on the y-axis. This technique has applications across a wide variety of studies. It is used in analysis of microtubule growth [1], kinetochore movement [2], lamellipodial advance or collapse[3] and probably most commonly, vesicle movement in neurons and other cell types[4]. Neuronal processes are particularly amenable to the use of kymograph analysis because of their inherent linear morphology. Their highly polarized structure of axons and dendrites provide built-in tracks along one axis where movement of a variety of biological structures can be followed. With fluorescent labeling, various processes and structures can be analyzed, such as trafficking of organelles[5,6], and other structures.

Kymographs contain a great deal of information about trafficking dynamics. Parameters available for analysis include the number of anterograde, retrograde, and stationary events, event speed, pause time, and event distance. If these parameters are to be measured under the condition where two proteins are trafficking together, events must be identified that coincide in both channels. A review of the literature shows that current kymograph analysis is limited to tracking one channel at a time[7–9]. The common method of creating kymographs from time dependent fluorescent microscope data is done on one channel producing one independent kymograph per marker with no direct connection between them, even though multiple channels can easily be acquired. To analyze two objects that co-traffic together, one would have to carefully examine the kymographs from both channels and decide which tracks are the same, or one could try to identify the coincident tracks from an overlay of the two channels. Tracks can merge and diverge, and such events are not easy to identify when looking at two separate images, so marking them can be a challenge. The whole process is laborious and time consuming. Detailed information from two kymographs would only be useful if all the data of interest are in one image. Thus far, there is no available open-source tool to generate one merged image from two kymographs. Our lab has been studying the dynamics of a variety of endosomal compartments and their inter-relationship in neurons using more than one endosomal marker [5,6,10]. The difficulty in finding an available open-source tool for co-localized trafficking analysis had led us to create such a program, which we are calling KymoMerge. KymoMerge addresses the issues discussed by automating the process of identifying co-located tracks in multi-channel kymographs and producing output that can be analyzed directly.

## Results and Discussion

### Program Description

The program we have created is written in Python and works as a plugin in ImageJ. The process is very straightforward. Kymographs need to be created for each channel. The available ImageJ plugin for creating kymographs works very well. The KymoMerge program then allows the user to set a threshold value for the individual kymograph images in each channel, which is determined as a percent of the peak intensity. The user can observe the original kymograph and the binary output and vary the threshold to obtain the best representation of the original data. Once the threshold is set, it is applied, and the image is converted to eight bits from the bit depth of the original image. This is then converted to a binary file and saved. Each pixel in the two channels is now set to a value of either 0 or 255. The program then goes pixel by pixel between the two kymographs comparing values and creating a new image. If the values are both zero, or one is zero and the other is 255 the new image is set at zero at that pixel. If they are both 255 (positive signal in both channels) then the value is set at 255. The result is an image consisting only of those pixels where positive signal is in both channels. This produces a kymograph containing tracks where the two proteins of interest are co-located, and the dynamics of the co-localized tracks can now be analyzed.

### KymoMerge versus Manual Kymograph Analysis

We created two sample data sets for analysis to test and validate the reliability and efficiency of the KymoMerge program. One consists of the neuronal membrane proteins, NSG1-cherry and NSG2-GFP. NSG1 and NSG2 are members of neuron-specific gene family of proteins. Both proteins are highly expressed in neurons and localized to a variety of endosomal compartments in dendrites (5). Our previous live imaging data showed that both proteins co-trafficked in neurons, and thus serve as a good example for analyzing the dynamics of two highly colocalized membrane cargos with low background noise signals. The other data set contains NSG1-cherry with GFP-Rab7, a late endosome marker. Rab7 is a small GTPase involved in regulating transport to late endocytic compartments. We have previously shown that Rab7 co-trafficked and regulated the endocytic transport of NSG1 in neurons (5). Like other Rabs, Rab7 is constantly cycling between the activated form (GTP bound) when it is recruited to membranes and inactivated form (GDP bound) after hydrolysis and when it is cytosolic. This feature allows us to test the ability of our program in extracting membrane-bound signals from cytosolic high background signals.

#### Visualizing co-localized tracks

In Figure 1 we give examples of typical output from KymoMerge. The images shown are from dendrite data where anterograde and retrograde movements are almost equally abundant and particularly hard to analyze. Figure 1a shows example kymographs from the Rab7, NSG1 data set. The top row shows the original 32-bit single channel images followed by an overlay of the two channels. NSG1 is in red and Rab7 is in green. Below are the binary outputs from KymoMerge. Figure 1b shows a similar example, in this case of the NSG1, NSG2 data set with NSG1 in red and NSG2 in green. Careful comparison of the original images in Figure 1 with the binary output of KymoMerge shows very similar tracks between the two. Due to the binary nature of the KymoMerge output it does not show intensity differences as in the original files, but this is not a parameter that is usually of interest in kymograph analysis. Intensity differences are only used initially in determining background levels and true signal during the thresholding stage before the binary output is created. Comparison of the overlayed and merged files also show similar characteristics. One of the principal strengths of our approach is that any track seen in the merged binary data is a co-located track. In contrast, it is difficult to discern coincident tracks in the original data.

**Figure 1.**
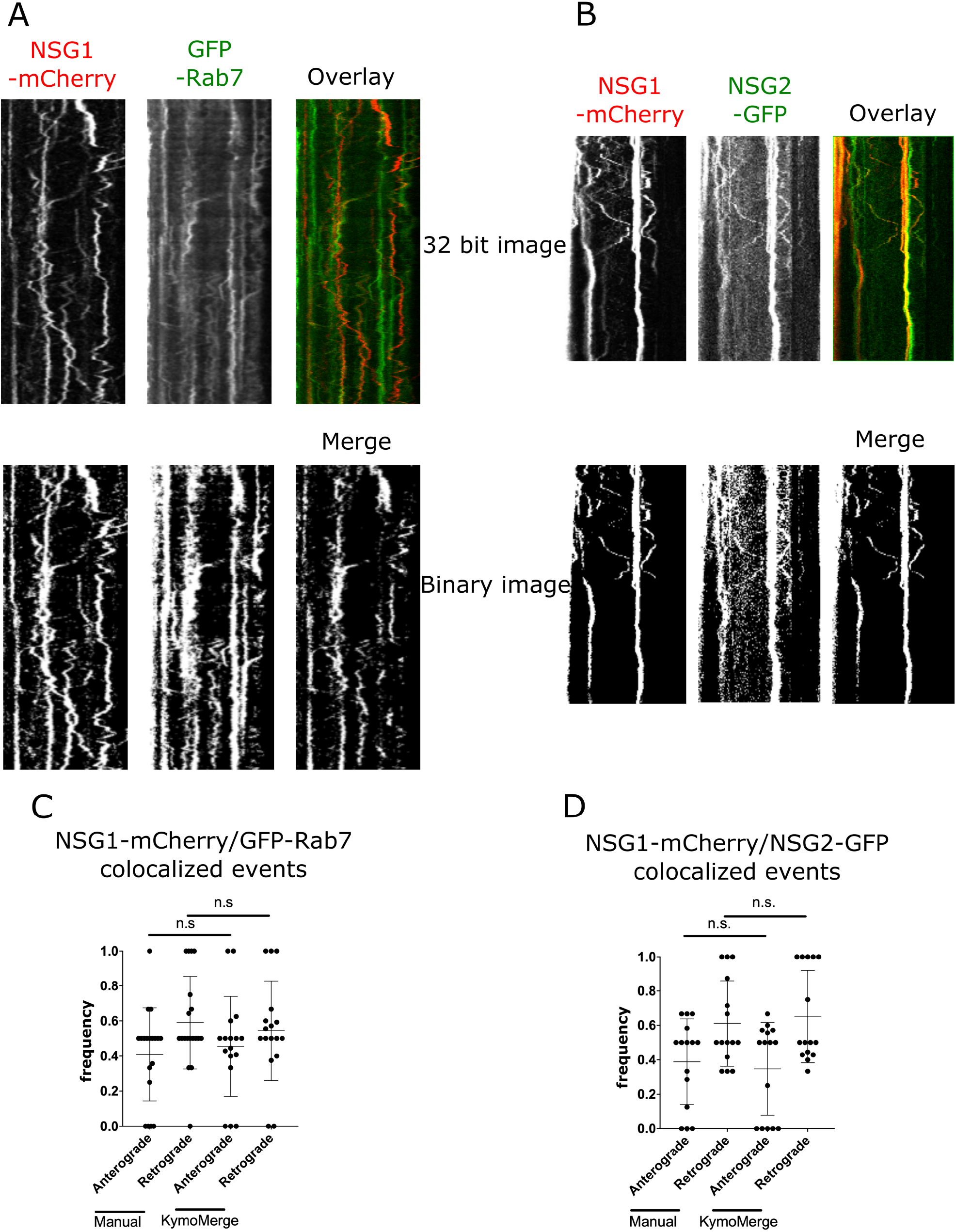
Comparison of of KymoMerge output and original data and consistency of results between them. **A & B**: Sample of source data of the NSG1-mCherry/GFP-Rab7 or NSG1-mCherry/NSG2-GFP data sets compared with the KymoMerge binary output. **C & D**: Comparison between manual counts of collocated tracks using original 32-bit images and KymoMerge created kymographs for the two data sets. N= 26 dendrites from 6 neurons in the NSG1-mCherry/Rab7-GFP data set. N= 24 dendrites from 7 neurons in the NSG1-mCherry/NSG2-GFP data set. Statistical results from Mann-Whitney test between anterograde or retrograde pairs under the two threshold conditions.

#### Testing the robustness of KymoMerge

Our approach is useful only if the program produces results that are robust and reliable, so we now address quality of output and potential issues. We thus determined whether KymoMerge produces results comparable to those from a careful count done by hand, as this is often how kymograph data is analyzed. For this purpose, we took the two data sets referenced in Figure 1a and 1b and independently counted anterograde and retrograde events manually, referencing either the original data and overlayed data as shown or using the merged binary output from KymoMerge. The results in Figure 1c and 1d show that there was no statistical difference between the two methods.

A fundamental problem in image analysis is distinguishing signal from noise. Cut-offs for noise vs signal needs to be set by the user and is thus often a more subjective decision. We addressed this issue by analyzing a data set with different threshold values to determine how much bias is introduced into the results through choice of threshold. Figure 2a shows examples of the image output from KymoMerge using a “high” or “low” threshold. The thresholds were set to represent a “reasonable” range that different experimenters might choose. The original data with the NSG1/Rab7 as an example is shown in Figure 2a along with the merged binary images created using different threshold values. The analysis was done by independently counting anterograde and retrograde events from the same data set after processing by KymoMerge. The results show that there is no statistically significant bias introduced through reasonably different thresholding values.

**Figure 2.**
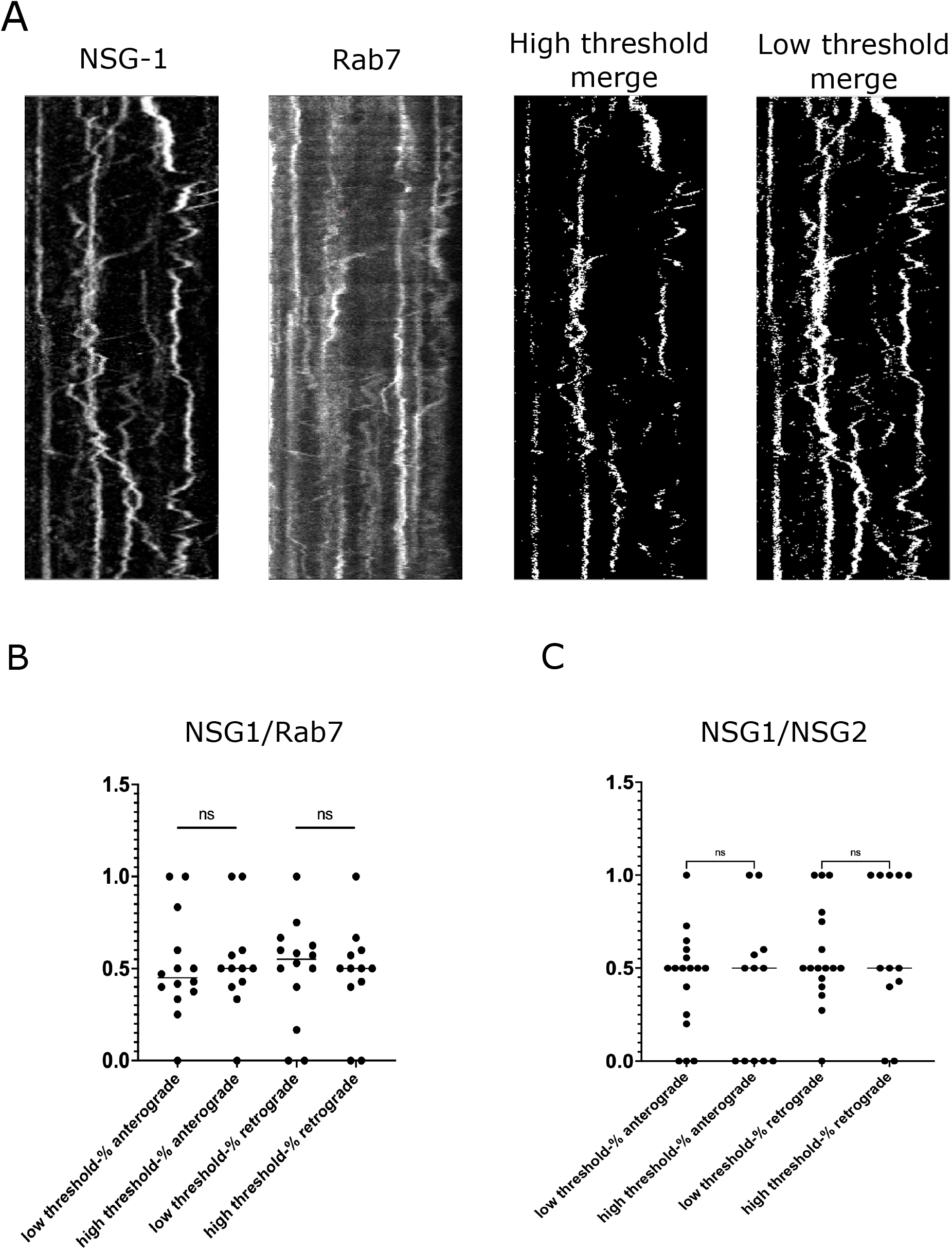
Example of threshold dependency of` KymoMerge output. **A**: Original 32-bit images from the NSG1-mCherry/Rab7-GFP data set and the merged two-channel binary output from KymoMerge with different threshold values. **B & C**: Anterograde and retrograde events were counted using KymoMerge colocalization output with low and high thresholds. For each kymograph the percent of anterograde and retrograde events are shown. N= 26 dendrites from 6 cells. Statistical results from Mann-Whitney test between anterograde or retrograde pairs under the two threshold conditions.

#### Limitations and Pitfalls

As discussed earlier, thresholding of images is often necessary in image analysis, but it is also highly subjective and, therefore, potentially problematic. With KymoMerge, thresholding requires attention particular to the process of creating binary images.

Issues can arise with kymographs that are particularly noisy. Figure 3a is an example of a very complicated and difficult to analyze image. The right half of the two original images have very little background and contains useful signal. The left halves have much more signal, and importantly for analysis, a high background, particularly in the NSG2 image. It is clear from the overlayed image that this region is impossible to interpret from the original data. The background makes all tracks appear co-located.

**Figure 3.**
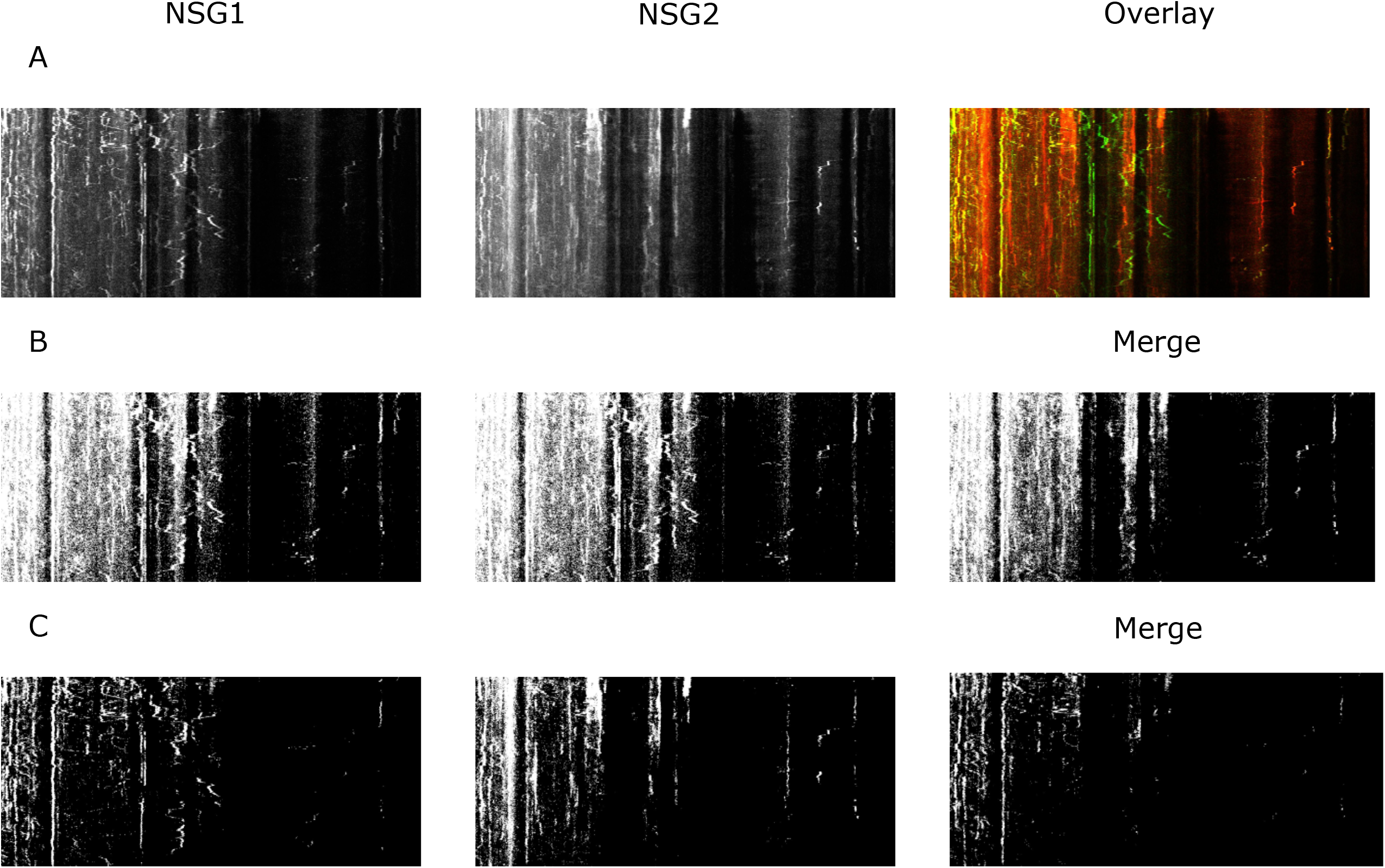
Examples of problems introduced through poor thresholding. **A:** Original 32-bit kymographs from NSG1-mCherry/NSG2-GFP data set with two-channel overlay. **B**: Binary KymoMerge output with low threshold at the level of. background. **C:** Binary output threshold set above background noise level.

A similar problem can occur with KymoMerge. Figure 3b shows the outputs that result from poor thresholding. This output is from an analysis of the NSG1-mCherry, NSG2-GFP data set with a threshold at a level that worked well for most images, but not well with this problematic one. There is a difficulty inherent in the process of creating the binary images. If one region is noisier than another, the noise can be of the same magnitude as the signal elsewhere. The result is the set threshold will either eliminate actual signal if it is at a level suitable for the noisier regions, or potentially background will be left creating false co-localized tracks. In the case of Figure 3B, with a too low threshold, the resulting noise produces a merged output that is uninterpretable. However, careful thresholding with KymoMerge can produce an accurate and interpretable merged image as shown in Figure 3C. Even though some signal is lost, what remains are true co-localized tracks that can be analyzed. For kymographs with areas of high background as in this example, it may be necessary to crop excessively noisy areas before analysis. Since these areas would generally be uninterpretable, analyzable data are not being lost and the rest of the image will produce accurate results. It is important to understand that any data set is only a sampling of the system being studied. It is better to err on the side of excluding false positive co-localized tracks and analyze only the resulting tracks that are truly co-localized. Because KymoMerge allows for individual screening and thresholding of the individual channels before the merged images are created any problematic kymographs can be identified by the user and properly processed.

## Conclusions

We have created a new tool that allows for the creation and analysis of co-localized two-channel kymographs. This method opens new experimental approaches to understanding vesicular transport. Comparisons can be made between overexpression of various proteins involved in transport, using either a control condition (GFP for instance) or differences between mutant variations of individual components. Images containing only tracks with both components can be directly analyzed. The approach could also be fairly easily expanded to allow for analysis of three or more co-trafficking markers. Additionally, there are methods of automated analysis that are only useful for single-channel data. Programs such as KymoAnalyzer[11] (for use with FIJI), Kymograph Clear[12], Kymograph Direct[12], or Kymobutler[13] can provide much richer data than manual analysis. First, however, single images need to be produced that contain only the multi-channel co-localized tracks. KymoMerge does that simply and accurately with maximal visual cross-check by the user.

## Methods

### Neuronal cultures and transfections

Neuronal cultures were prepared from E18 rat hippocampi as described [5]. All experiments were performed in accordance with Institutional Animal Care and Use Committee guidelines and regulations. Rats were euthanized by CO2 inhalation in a chamber. CO2 flow was maintained until the rat was deeply anesthetized (verified via toe-pinch) and in CO2 narcosis. Death was confirmed by decapitation and embryos removed.

Hippocampi from all pups in one litter were combined and thus contained male and female animals. Cells were plated on poly-l-lysine–coated coverslips and incubated with plating medium containing DMEM with 10% horse serum. For live-imaging use, neurons were plated on a 35-mm glass-bottomed microwell dish (MatTek). After 4 h, the plating medium was removed and replaced with serum-free medium supplemented with B27 (Thermo Fisher Scientific), and neurons were cultured for 7–10 DIV for experimental use. Transfections were performed with Lipofectamine 2000 (Invitrogen).

### Live imaging and kymograph analysis

Live imaging was performed as described by [5] In brief, neurons at DIV 7 were transfected for 36–40 h with the following plasmid combinations: *NSG1-mCherry* with *GFP-Rab7* or *NSG1-mCherry* with *NSG2-GFP*. Prolong Live Antifade (<u>P36974</u>; Invitrogen) was added right before live imaging. All live imaging was performed on a 37°C heated stage in a chamber with 5% CO2. All dual live imaging was conducted on an inverted LSM880 confocal microscope using a 40× water objective (LD-C Apochromat 1.2W). Images from dual channels were acquired simultaneously with bidirectional scan-frame mode every second for 300–500 frames and with tight gate settings to reduce any overlapping fluor spectra. Laser lines at 488 nm for GFP and at 594 nm for mCherry expression were used. Kymographs were generated using FIJI (ImageJ; National Institutes of Health). For the manual counting of colocalized events, images from each channel were overlaid and collocated retrograde and anterograde events ≥ 2μm were counted.

### Comparison algorithm

The comparison algorithm takes as input a pair of kymographs to be compared. We can think of these as n by m matrixes of pixels, with pixels valued in P = {0, 1, 2, …255}, according to intensity. The algorithm starts with pre-processing procedure.

Given kymograph x ∈ M_w,h_(P), pre-processing is performed as follows:

I. Thresholding: A value t ∈ (0, 100) is inputted as threshold. We use t to calculate T (x), given as T (x) := t/100 ∗ max_xi,j_ ∈x |x_i,j_ |, or in other words, t% of the maximum activation exhibited by kymograph x. Threshold value t is set per individual kymograph, at users discretion.
II. Binarization: Given pixel x_i,j_, the binary of that pixel, denoted B(x_i,j_), is given as

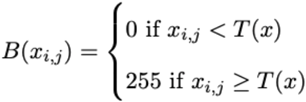
III. Merging: Takes as input kymographs x,y and returns merged kymograph, given as

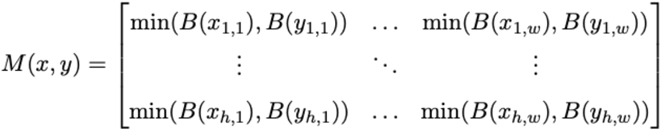

Returns grid of minimum values of binaries for each corresponding pixel pair in kymographs x and y. This is used to determine sites where both of kymographs exhibited activation above the threshold value.

### Using KymoMerge

KymoMerge is available to download from Github at: https://github.com/alduston/kymomerge.

Using KymoMerge relies on two principal components: the KymoMerge.py script and the data directory. To begin using the merge tool open up an instance of FIJI and open the KymoMerge.py script in the FIJI console. The program can also be installed as a plugin as described in the ImageJ support pages.

Input directory 1 should be a directory containing a series of named tif files, using the naming convention ‘filename’-’i’.tif where i is the given tif files ‘index’ in the folder. For instance, given 3 files in ‘group A’ directory, using ‘kymo’ as file name, the tif files would be given {kymo-1.tif, kymo-2.tif, kymo-3.tif}.

Input directory 2 should be a directory containing a series of named tif files, using the naming convention ‘filename’-’i’.tif where i is the given tif files ‘index’ in the folder. This index and file name should correspond to the index of input directory 1.The algorithm will compare files with corresponding indexes. For instance, given 3 files in ‘groupB’ directory, using ‘kymo’ as the file name as before, the .tif files would be given {kymo-1.tif, kymo-2.tif, kymo-3.tif}.

Threshold 1 should be the threshold value t1 ∈ (0, 100) appropriate to the kymographs input directory 1. This will be adjusted for each file by the user as the program runs to the optimum value for each image.

Threshold 2 Should be the threshold value t2 ∈ (0, 100) appropriate to the kymographs in input directory 2.

After inputting above values appropriate for individual kymographs, the algorithm will run over the data and save the outputs into a directory named (using names from the examples above), ‘groupA-groupB output’, located where the input directories are located. This directory will contain a sub-directory containing the outputs for given tif file pair, named as ‘filename-i ‘, or ‘kymo-i ‘in the example case. Each of these sub-directories will contain 3 output .tif files.:

I. ‘groupA’ ‘filename’-’i’.tif : Binarized thresholded version of i’th file in groupA directory.
II. ‘groupB’ ‘filename’-’i’.tif : See I, for group B
III. ‘groupA’-’groupB’ collocated ‘filename’-’i’.tif : Result of merging I & II. Corresponds to sites where A and B are activated beyond threshold level. Ex: Directory kymo-1 in groupA-groupB output directory is given {groupA kymo-1.tif, groupB kymo-1.tif, groupA-groupB collocated kymo-1.tif} In actual use when the program is opened the following dialog box will appear:

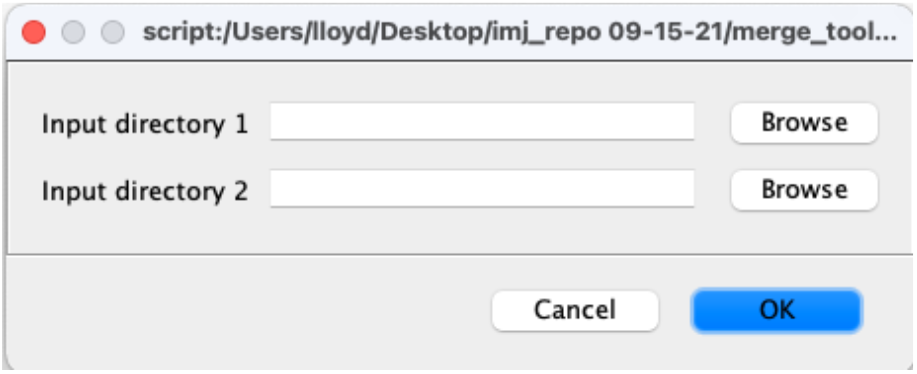

After choosing the directories containing the kymographs for the two channels there are two following steps.

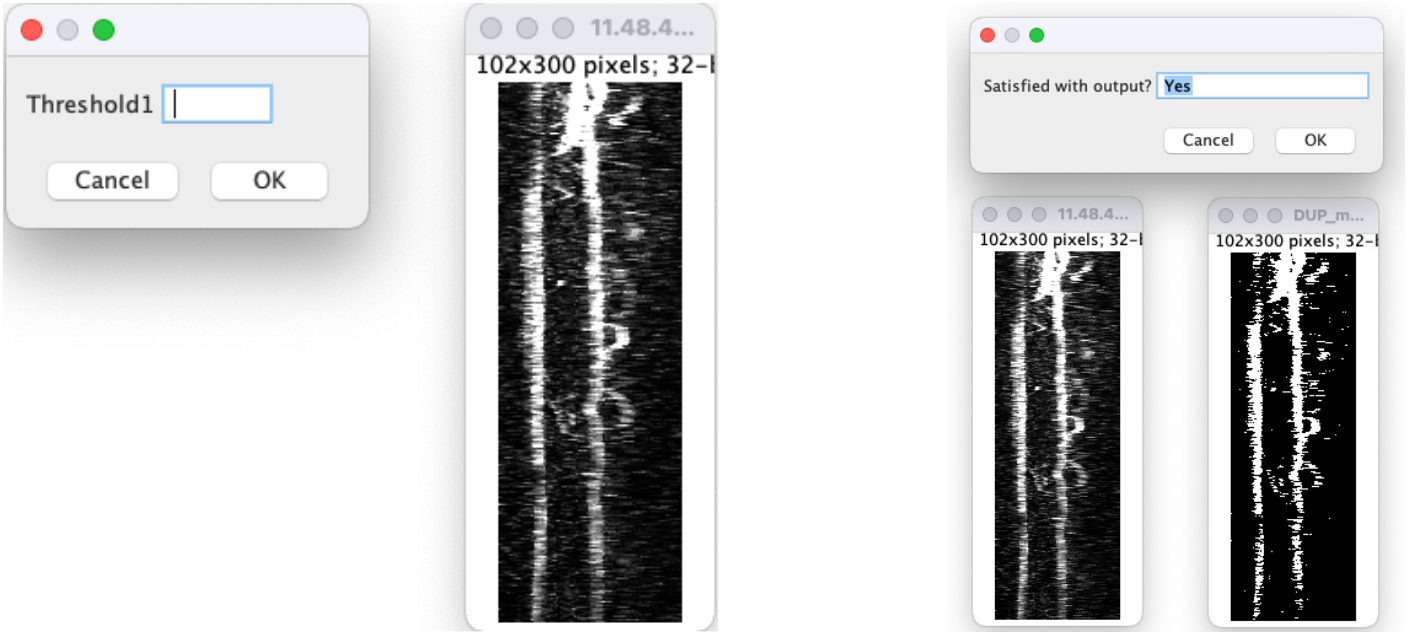

The dialog box and image on the left comes up and shows the original kymograph along with an input to set the threshold. The contrast for the original image is automatically increased so the data are clearly visible. When a value is chosen for the threshold, the binarized image appears and can be compared to the original as shown with the dialog box and images on the right. At this point the user can accept the value by typing in yes or choosing OK or try a new value by typing in no or choosing Cancel. Once a value is accepted the next image opens. Once all images are thresholded, they will be processed and the output (individual binary images and merged image) will be available for further analysis as described above. Should the user want to quit the analysis at any point simply type “quit” into the dialog box.

